# Liver Fibroblast Growth Factor 21 (FGF21) is Required for the Full Anorectic Effect of the Glucagon-Like Peptide-1 Receptor Agonist Liraglutide in Male Mice fed High Carbohydrate Diets

**DOI:** 10.1101/2023.01.03.522509

**Authors:** Thao D. V. Le, Payam Fathi, Amanda B. Watters, Blair J. Ellis, Nadejda Bozadjieva-Kramer, Misty B. Perez, Andrew I. Sullivan, Jesse P. Rose, Laurie L. Baggio, Jacqueline Koehler, Jennifer L. Brown, Michelle B. Bales, Kaitlyn G. Nwaba, Jonathan E. Campbell, Daniel J. Drucker, Matthew J. Potthoff, Randy J. Seeley, Julio E. Ayala

## Abstract

Glucagon-like peptide-1 receptor (GLP-1R) agonists and fibroblast growth factor 21 (FGF21) confer similar metabolic benefits. Studies report that GLP-1RA induce FGF21. Here, we investigated the mechanisms engaged by the GLP-1R agonist liraglutide to increase FGF21 levels and the metabolic relevance of liraglutide-induced FGF21. We show that liraglutide increases FGF21 levels via neuronal GLP-1R activation. We also demonstrate that lack of liver *Fgf21* expression confers partial resistance to liraglutide-induced weight loss. Since FGF21 reduces carbohydrate intake, we tested whether the contribution of FGF21 to liraglutide-induced weight loss is dependent on dietary carbohydrate content. In control and liver *Fgf21* knockout (Liv^*Fgf21*-/-^) mice fed calorically matched diets with low- (LC) or high-carbohydrate (HC) content, we found that only HC-fed Liv^*Fgf21*-/-^ mice were resistant to liraglutide-induced weight loss. Similarly, liraglutide-induced weight loss was partially impaired in Liv^*Fgf21*-/-^ mice fed a high-fat, high-sugar (HFHS) diet. Lastly, we show that loss of neuronal β-klotho expression also diminishes liraglutide-induced weight loss in mice fed a HC or HFHS diet, indicating that FGF21 mediates liraglutide-induced weight loss via neuronal FGF21 action. Our findings support a novel role for a GLP-1R-FGF21 axis in regulating body weight in the presence of high dietary carbohydrate content.

## Introduction

Obesity is one of the largest health challenges in recent decades. Nearly 1 in 3 U.S. adults are overweight^1^ and more than 2 in 5 have obesity^1,2^. In addition to their direct action in the pancreas to stimulate insulin secretion, glucagon-like peptide-1 (GLP-1) receptor (GLP-1R) agonists (GLP-1RA) comprise a class of drugs that also promote weight loss^3–9^. This weight loss effect is primarily due to a reduction in food intake resulting from GLP-1RA acting on several regions of the brain^10–14^. However, the mechanisms by which brain GLP-1R activation promotes weight loss remain unclear.

Fibroblast growth factor 21 (FGF21) is a hormone mainly produced by the liver in response to metabolic challenges including low protein and high carbohydrate consumption^15–20^. Interestingly, FGF21 displays effects that overlap with many of those associated with GLP-1R activation such as improved glycemic control^21–24^, weight reduction^21,25–27^, and suppression of carbohydrate intake^28,29^. A connection between GLP-1RA and FGF21 has been implicated in studies showing that GLP-1R activation induces FGF21 production^30–35^. However, since GLP-1RA reduce caloric intake, and reduced caloric intake is a key stimulus for FGF21 production^36–38^, it remains unclear whether GLP-1RA-induced stimulation of FGF21 production is secondary to its food intake-suppressing properties. Furthermore, the contribution of FGF21 to the metabolic benefits of GLP-1RA has not been thoroughly addressed. Given the clinical benefits of GLP-1RA and FGF21, the therapeutic implications of a GLP-1 R-FGF21 axis merit investigation.

In the current study, we demonstrate that the therapeutic GLP-1RA liraglutide acts on neuronal GLP-1R to increase circulating FGF21 levels in a food intake-independent manner. We also show that liraglutide-induced FGF21 is required for the full weight-lowering effects of GLP-1R activation in mice fed a chow-diet. Given the reported role of FGF21 as a feedback inhibitor of carbohydrate intake^28,29^, we hypothesized that FGF21 plays a specific role to reduce the intake of carbohydrates in response to GLP-1RA treatment. We support this hypothesis by showing that mice lacking liver *Fgf21 (Liv^Fgf21-/-^*) are resistant to the weight-lowering effect of liraglutide only when fed high-carbohydrate diets and not when fed low-carbohydrate diets. Lastly, we show that central FGF21 signaling is required for FGF21 to mediate the weight loss action of liraglutide. Taken together, these findings suggest that the weight loss response to liraglutide is influenced by carbohydrate content in a FGF21-dependent manner. This may have significant clinical implications because it suggests that some of the observed variability in weight loss efficacy of GLP-1RA^39–41^ may be influenced by the diets consumed by and/or *Fgf21* polymorphisms present in people taking these drugs.

## Results

### GLP-1RA increase FGF21 independently of its food intake-suppressing effects

Studies show that GLP-1RA induce FGF21 levels in several mouse models^30–35^. However, since fasting or a state of nutrient deficit is a major stimulus for FGF21 production^36–38^, it remains unclear whether the increase in FGF21 following GLP-1R agonist treatment is due to the food intake suppressive effects of these compounds. We first tested whether the GLP-1RA liraglutide increased FGF21 levels independently of its effects on food intake by administering vehicle or liraglutide to 4 hour (h)-fasted, male C57BL/6J mice and measuring plasma FGF21 levels at 0 and 7 h following treatment. All mice remained without food for the duration of the study period (**Figure 1A**). Plasma FGF21 levels were significantly higher 7 h following administration of liraglutide (**Figure 1B**) compared to vehicle-treated controls. A similar effect was observed in mice treated with the GLP-1RA exendin-4 (**Supplemental Figure 1**). Elevated circulating FGF21 levels in response to liraglutide were associated with a significant increase in *Fgf21* mRNA in the liver (**Figure 1C**), consistent with previous studies showing that circulating FGF21 is predominantly secreted from this organ^38^. To further control for the food intake and subsequent weight-lowering effect of liraglutide, we treated *ad libitum-fed* male C57BL/6J mice with vehicle or liraglutide for 2 days and added a third group of mice pair-fed to weight match the liraglutide-treated group (**Figure 1D** and **1E**). Circulating FGF21 levels were significantly higher in liraglutide-treated mice compared to vehicle-treated mice and mice pair-fed to weight match the liraglutide-treated group (**Figure 1F**). These results demonstrate that GLP-1RA increase FGF21 independently of either their food intake-suppressive effects or their ability to promote weight loss.

**Figure 1.**
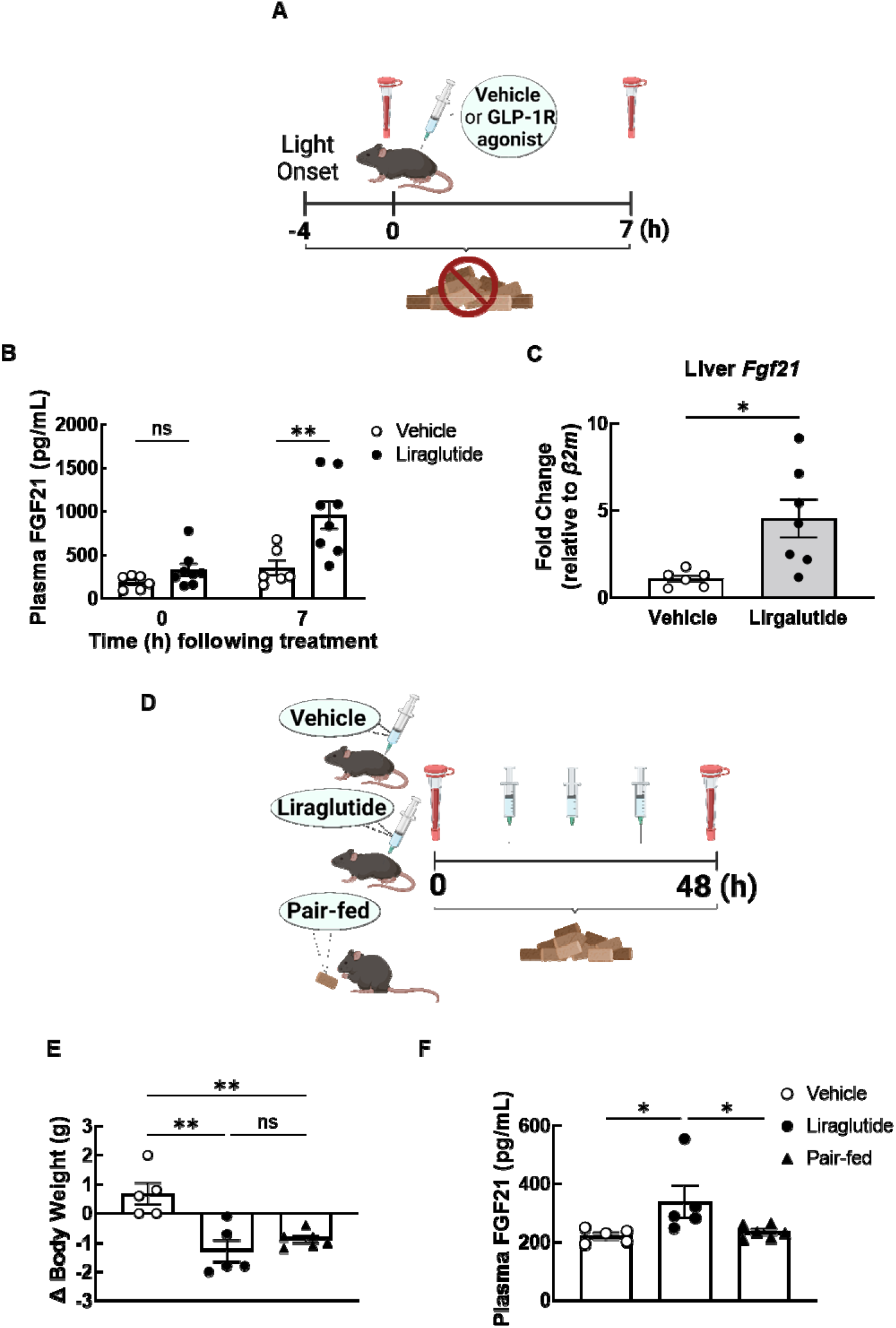
GLP-1RA stimulate FGF21 independent of their effect on food intake. (**A**) Study outline for B-C. (**B**) Plasma FGF21 levels at 0 and 7 h following treatment in fasted C57BL6/J male mice treated with vehicle or liraglutide (400 μg/kg) (Mixed-effects analysis: Interaction, F (1, 12) = 6.62, P = 0.0244; N = 6-8). (**C**) Relative change in liver *Fgf21* gene expression in fasted C57BL6/J male mice 7 h following treatment with vehicle or liraglutide (400 μg/kg) (Unpaired t-test: P = 0.0152; N = 6-8). (**D**) Study outline for E-F. (**E-F**) Changes in body weight (E; One-way ANOVA: Interaction, F (2, 13) = 12.02, P = 0.0011; N = 5-6)) and plasma FGF21 levels (F; One-way ANOVA: Interaction, F (2, 13) = 4.111, P = 0.0414; N = 5-6) in C57BL6/J male mice *ad libitum-fed* and treated with vehicle or liraglutide (200 μg/kg, *b.i.d*), or pair-fed to weight match the liraglutide-treated group for 48 hours. Data are shown as mean ± SEM, ns not significant, * P < 0.05, ** < 0.01.

### Central nervous system GLP-1R and liver PPARα are required for liraglutide to increase plasma FGF21

While plasma FGF21 is primarily derived from the liver^38^, the GLP-1R is not expressed in hepatocytes^42–44^. We, therefore, tested whether neuronal or pancreatic β-cell GLP-1R expression is required for liraglutide to increase FGF21 levels. Using the same protocol as in Figure 1A, we administered vehicle or liraglutide to fasted control mice and mice lacking the GLP-1R in neurons targeted by the Wnt1-*Cre2* driver (Wnt1^*Glp1r-/-*^) and also in mice lacking the GLP-1R specifically in glutamatergic neurons (Vglut2^*Glp1r-/-*^). We used these models since both are resistant to the weight lowering effects of liraglutide^13,14^. Liraglutide failed to induce FGF21 in both Wnt1^*Glp1r-/-*^ (**Supplemental Figure 2A** and **2B**) and Vglut2^*Glp1r-/-*^ (**Figure 2A** and **2B**) mice. In contrast, the stimulatory effect of liraglutide on FGF21 remained intact in mice lacking the GLP-1R in pancreatic β-cells (**Fig. 2C** and **2D**), another major site of GLP-1R expression and actions^44^. Taken together, these findings suggest that that neuronal GLP-1R, and not β-cell GLP-1R, is required for liraglutide to increase FGF21.

**Figure 2.**
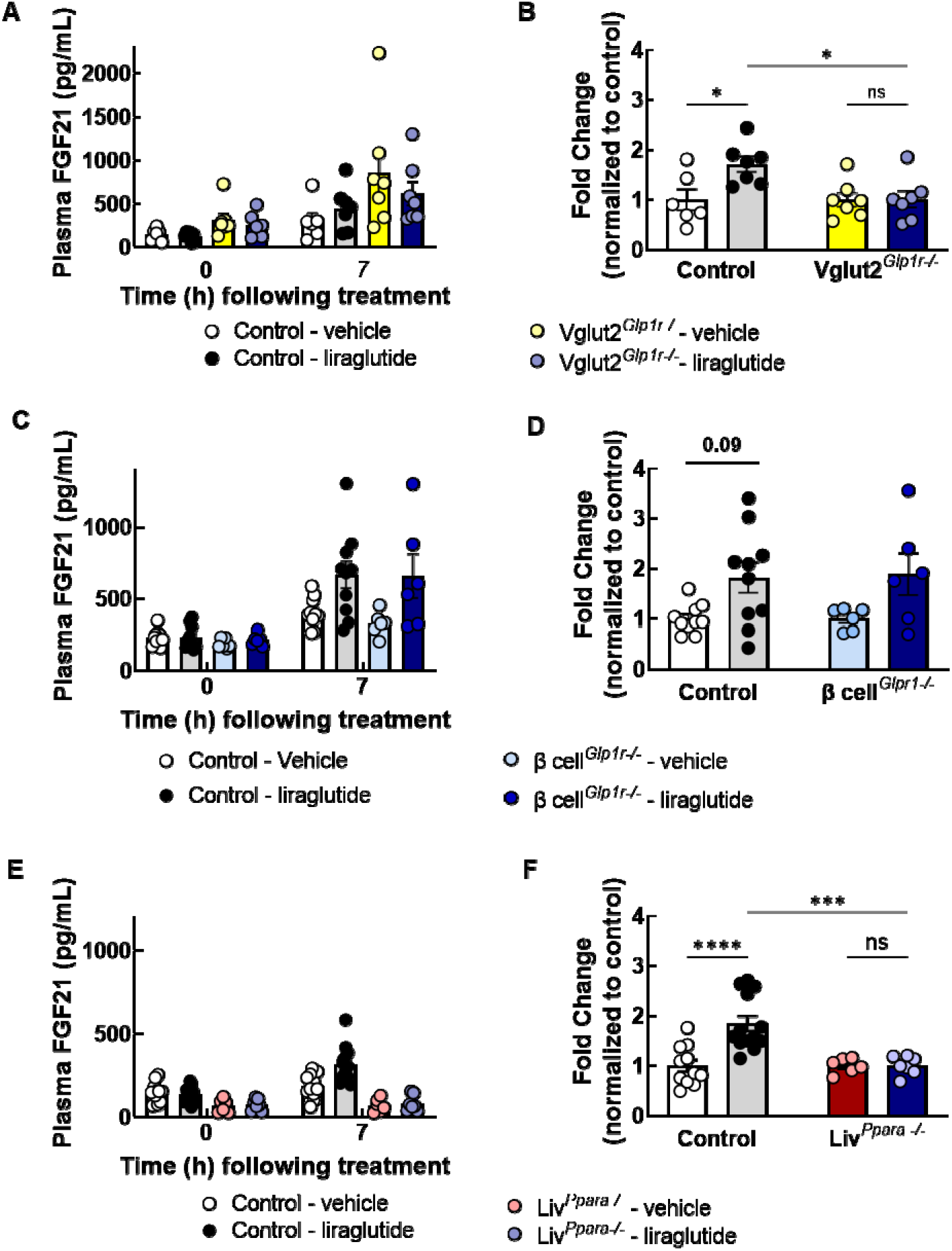
Central GLP-1R and liver PPARα are required for liraglutide to stimulate plasma FGF21. Absolute (**A**, C, E) and relative change (B, D, F) in FGF21 levels at 0 and 7 h post treatment with vehicle or liraglutide (400 μg/kg) in fasted control and respective knockout mice. (**A-B**) Control and vGLUT2+ neuron-*Glp1r* knockout (Vglut2^*Glp1r-/-*^) mice (A, Mixed-effects analysis: Interaction, F (1, 23) = 2.021, P = 0.1686; Genotype Effect, F (1, 23) = 5.965, P = 0.0227; B, Mixed-effects analysis: Interaction, F (1, 23) = 4.571, P = 0.0434; N = 5-6). (**C-D**) Control and β cell-*Glp1r* knockout (β cell^*Glp1r-/-*^) mice (C, Mixed-effects analysis: Interaction, F (1, 56) = 0.05070, P = 0.8227; Genotype Effect, F (1, 56) = 11.28, P = 0.0014; D, Mixed-effects analysis: Interaction, F (1, 28) = 0.01663, P = 0.8983; Genotype Effect, F (1, 28) = 10.41, P = 0.0032; N = 6-10). (**E-F**) Control and liver *Ppara* knockout (Liv^*Ppara-/-*^) mice (E, Mixed-effects analysis: Interaction, F (1, 66) = 9.627, P=0.0039; Liraglutide-treated control vs. Liv^*Ppara-/-*^ at 7 h, P < 0.0001; **F**, Mixed-effects analysis: Interaction, F (1, 33) = 8.775, P = 0.0056; N = 6-13). Data are shown as mean ± SEM, ns not significant, * P < 0.05, *** P < 0.001, and **** P < 0.0001.

To verify that liraglutide-induced circulating FGF21 originates from the liver, we administered vehicle and liraglutide to fasted control mice and mice lacking liver PPARα, a key regulator of liver FGF21 production particularly in response to fasting. Plasma FGF21 levels were increased in liraglutide-treated control mice but not in liraglutide-treated liver *Ppara* knockout (Liv^*Ppara*-/-^) mice (**Figure 2E** and **2F**). Taken together, these findings suggest that liraglutide engages neuronal GLP-1R to induce FGF21 production, and increased FGF21 production requires liver PPARα expression.

### FGF21 is partially required for liraglutide-induced weight loss

To investigate the contribution of FGF21 to the weight-lowering effects of GLP-1R activation, we chronically administered vehicle or liraglutide to chow-fed control and liver *Fgf21* knockout (*Liv^Fgf21-/-^*) mice. Circulating FGF21 levels were almost undetectable in both vehicle- and liraglutide-treated Liv^*Fgf21*-/-^ mice (**Supplemental Figure 3A**). Unlike acute treatments, chronic liraglutide dosing did not significantly elevate circulating FGF21 levels in chow-fed mice. Nevertheless, chow-fed Liv^*Fgf21*-/-^ mice were partially resistant to liraglutide-induced weight loss (**Figure 3A** and **3B**). These experiments were conducted in mice housed in metabolic cages to continuously measure metabolic parameters such as food intake and energy expenditure during chronic liraglutide treatment. The partial resistance to liraglutide-induced weight loss in *Liv^Fgf21-/-^* mice was due, in part, to an attenuated reduction in food intake, particularly during the first day of treatment (**Figure 3C**). The average daily caloric intake and total calories consumed throughout the liraglutide treatment period were reduced to a lesser degree in *Liv^Fgf21-/-^* mice (**Figure 3D** and **Supplemental Figure 3B**). Liraglutide treatment reduced energy expenditure (EE) in both control and Liv^*Fgf21*-/-^ mice, but this effect was more pronounced in liraglutide-treated Liv^*Fgf21*-/-^ mice (**Figures 3E**, **3F**, and **Supplemental Figure 3C**). The significant body weight difference between liraglutide-treated control and Liv^*Fgf21*-/-^ mice (**Figure 3G**) was associated with an attenuated, albeit not significant, reduction in fat-free (i.e., lean) mass in Liv^*Fgf21*-/-^ mice (**Figure 3H**). Changes in fat mass did not differ between liraglutide-treated control and Liv^*Fgf21*-/-^ mice (**Figure 3I**).

**Figure 3.**
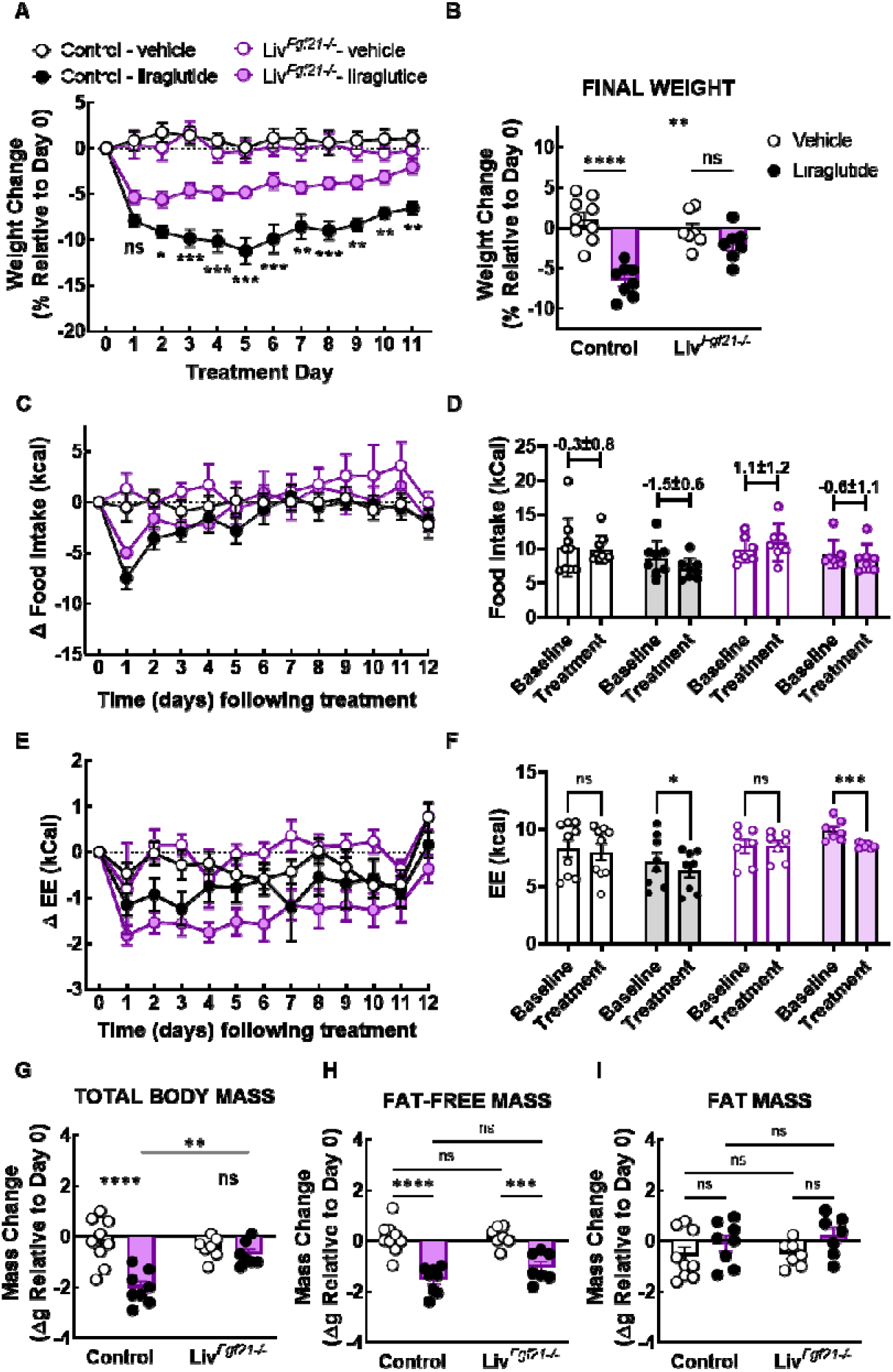
Liver FGF21 mediates the appetite- and weight-lowering actions of liraglutide. Chow-fed control and liver *Fgf21* knockout (Liv^*Fgf21-/-*^) mice housed in metabolic cages and treated with vehicle or liraglutide (200 μg/kg, *b.i.d*) for 12 days (N = 7-9). (**A-B**) Relative weight loss over time (A; Mixed-effects analysis: Interaction, F (11, 156) = 1.920, P = 0.0405) and relative weight loss on the last day of treatment (B; Mixed-effects analysis: Interaction, F (1, 27) = 13.11 P = 0.0012). (**C**) Change in food intake relative to baseline (Mixed-effects analysis: Interaction, F (36, 324) = 2.154, P = 0.0003; Genotype Effect, F (3, 27) = 1.271, P = 0.3042). (**D**) Average food intake at Baseline (Day 0) and during the Treatment period (average of Day 1-12) (Mixed-effects analysis: Interaction, F (3, 27) = 1.271, P = 0.3042; Genotype Effect, F (3, 27) = 2.137, P = 0.1189). (**E**) Change in energy expenditure (EE) relative to baseline (Mixed-effects analysis: Interaction, F (36, 324) = 1.151, P = 0.2606; Genotype Effect, F (3, 27) = 5.035, P = 0.0067). (**F**) Average food intake at Baseline (Day 0) and during the Treatment period (average of Day 1-12) (Mixed-effects analysis: Interaction, F (3, 27) = 5.035, P = 0.0067; Genotype Effect, F (3, 27) = 2.615, P = 0.0716). (**G-I**) Total body mass (G; Mixed-effects analysis: Interaction, F (1, 27) = 10.60, P = 0.0030), fat-free (or lean) mass (H; Mixed-effects analysis: Interaction, F (1, 27) = 1.383, P = 0.2499; Genotype Effect, F (1, 27) = 1.983, P = 0.1705), and fat mass **(I;** Mixed-effects analysis: Interaction, F (1, 27) = 0.2012, P = 0.6573; Genotype Effect, F (1, 27) = 0.4077, P = 0.5285) at the end of treatment. Data are shown as mean ± SEM, ns not significant, * P < 0.05, ** P < 0.01, *** P < 0.001, and **** P < 0.0001 for comparisons between liraglutide-treated control vs. *Liv^Fgf21-/-^* mice (A, C, E) and those delineated by lines (B, F, G-I).

### FGF21 mediates liraglutide-induced weight loss specifically in the context of high-carbohydrate diets

Since FGF21 suppresses carbohydrate intake and sweet preference in rodents^28,29^ and has been associated with these phenotypes in humans^45–50^, we hypothesized that FGF21 contributes specifically to a reduction in body weight by liraglutide in the presence of high-carbohydrate diets. To test this hypothesis, we placed control and Liv^*Fgf21*-/-^ mice on calorically matched, low-fat diets with either low (LC) or high carbohydrate (HC) content (30%, LC and 70%, HC, respectively) for 4 weeks followed by a 2-week treatment with vehicle or liraglutide while mice remained on the respective diet. Regardless of diet and treatment, circulating FGF21 levels were almost undetectable in Liv^*Fgf21*-/-^ mice (**Supplemental Figure 4A** and **4B**). Liraglutide reduced body weight to a greater degree in HC-fed control mice (**Figure 4C** and **4D**) than in LC-fed control mice (**Figure 4A** and **4B**). Importantly, only HC-fed Liv^*Fgf21*-/-^ mice were significantly resistant to liraglutide-induced weight loss (**Figure 4C** and **4D**), suggesting that FGF21 contributes to the weight loss effects of liraglutide in mice maintained on low-fat, high-carbohydrate diets. Next, we examined the relevance of these findings in the context of a high-fat, high-sugar (HFHS) diet, which better resembles a typical American diet. We placed cohorts of control and Liv^*Fgf21*-/-^ mice on a diet containing 45% fat and 35% carbohydrate, 50% of which is sucrose, for 1 or 4 weeks followed by treatment with vehicle or liraglutide for 2 weeks. It is important to note that mice did not gain weight when fed the HFHS diet for 1 week. The 1-week HFHS cohort is therefore included to control for any potential effects of the diet-induced weight gain observed in mice fed the same diet for 4 weeks. As seen in **Figures 4E** - **4H**, when maintained on either 4 weeks (**Figure 4E** and **4F**) or 1 week (**Figure 4G** and **4H**) of HFHS diet prior to treatment, Liv^*Fgf21*-/-^ mice lost less weight than their control counterparts when dosed with liraglutide. Taken together, these results support our hypothesis that FGF21 mediates the weight lowering actions of liraglutide specifically in the context of carbohydrate/sugar-rich diets.

**Figure 4.**
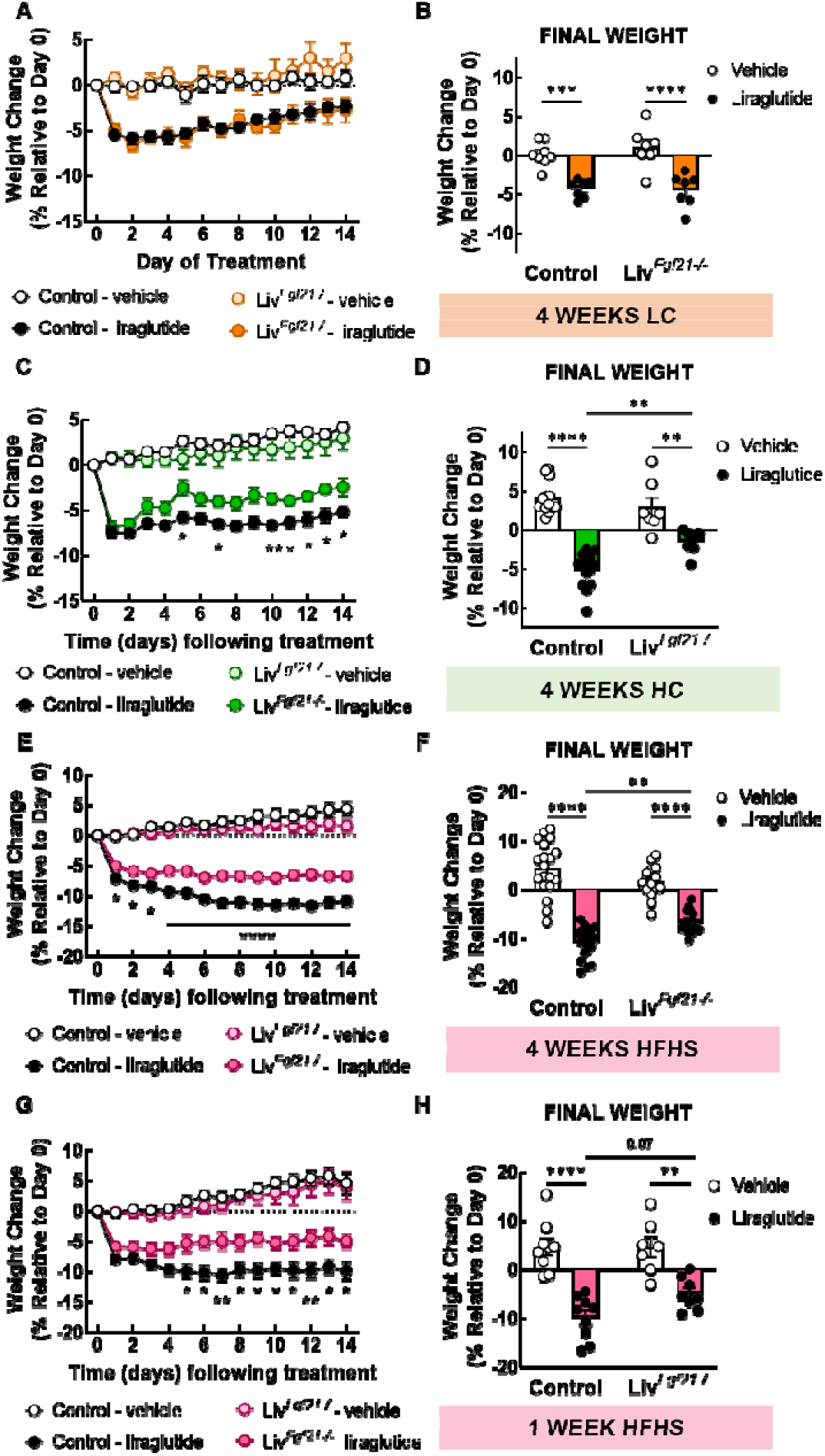
Liver FGF21 contributes to liraglutide-induced weight loss in the context of high-carbohydrate diets. Control and liver *Fgf21* knockout (Liv^*Fgf21-/-*^) mice fed a low- or high-carbohydrate diet for 4 weeks (A-F) or 1 week (G-H) followed by treatment with vehicle or liraglutide (200 μg/kg, *b.i.d*) for 14 days. (**A-B**) Relative weight loss over time (A; Mixed-effects analysis: Interactions, F (14, 168) = 0.5293, P = 0.9134; Genotype Effect, F (1, 12) **=** 0.02792, P = 0.8701; N = 7-8) and on the last day of treatment (B; Mixed-effects analysis: Interactions, F (1, 25) = 0.5317, P = 0.4727; N = 7-8) of mice fed a low-fat, low-carbohydrate diet for 4 weeks. (**C-D**) Relative weight loss over time (C; Mixed-effects analysis: Interactions, F (14, 266) = 1.865, P = 0.0303; N =7-14) and on the last day of treatment (D; Mixed-effects analysis: Interactions, F (1, 37) = 10.35, P = 0.0027; N = 7-14) of mice fed a low-fat, high-carbohydrate diet for 4 weeks. (**E-F**) Relative weight loss over time (E; Mixed-effects analysis: Interactions, F (14, 434) = 7.323, P < 0.0001; N = 14-19) and on the last day of treatment (F; Mixed-effects analysis: Interactions, F (1, 62) = 13.12, P = 0.0006; N = 14-19) of mice fed a high-fat, high-sugar diet for 4 weeks. (**G-H**) Relative weight loss over time (G; Mixed-effects analysis: Interactions, F (14, 210) = 4.004, P < 0.0001; N = 7-9) and on the last day of treatment (H; Mixed-effects analysis: Interactions, F (1, 29) = 2.291, P = 0.1410; Genotype Effect, F (1, 29) = 2.303, P = 0.1400; N = 7-9) of mice fed a high-fat, high-sugar diet for 1 week. Data are shown as mean ± SEM, ns not significant, * P < 0.05, ** P < 0.01, *** P < 0.001, and **** P < 0.0001 for comparisons between liraglutide-treated control vs. Liv^*Fgf21-/-*^ mice (C, E, G) and those delineated by lines (D, F, H).

### FGF21 mediates liraglutide-induced weight loss via its actions in the central nervous system

We next examined the target engaged by FGF21 to facilitate the weight-lowering effects of liraglutide. FGF21 signals to a receptor complex comprised of FGF receptor 1c (FGFR1c) and its co-receptor, β-klotho (KLB)^51–53^. FGF21 binds directly to KLB, leading to subsequent binding to FGFR1c and triggering downstream signaling events. As FGFR1c is ubiquitously expressed, KLB expression confers tissue specificity for FGF21^54^. Previous studies show that obese mice lacking *Klb* expression in the forebrain (Camk2a^*Klb-/-*^) but not those lacking *Klb* expression in the hindbrain (Phox2b^*Klb-/-*^) are resistant to the suppressive effects of recombinant FGF21 on body weight and energy expenditure^55^. More recent studies showed that mice lacking *Klb* expression in glutamatergic neurons (Vglut2^*Klb-/-*^), and not those lacking *Klb* expression in GABAergic or dopaminergic neurons, were resistant to FGF21-induced reduction in sugar intake and sweet preference^56^. We, therefore, tested the hypothesis that FGF21 acts in these neuronal populations to facilitate the weight loss induced by liraglutide. When placed on the same HC diet as that used in **Figures 4C** and **4D**, liraglutide-treated Camk2a^*Klb-/-*^ mice displayed partial resistance to weight loss compared to liraglutide-treated control mice (**Figure 5A** and **5B**). Similarly, Vglut2^*Klb-/-*^ mice maintained on 1 week of the same HFHS diet used in **Figure 4G** and **4H** were also resistant to the weight-lowering effects of liraglutide compared to control mice (**Figure 5C** and **5D**). These findings suggest that central FGF21 signaling is required for the full effect of liraglutide-induced weight loss, supporting our hypothesis that liraglutide-induced FGF21 signals to the central nervous system to facilitate the weight-lowering actions of liraglutide.

**Figure 5.**
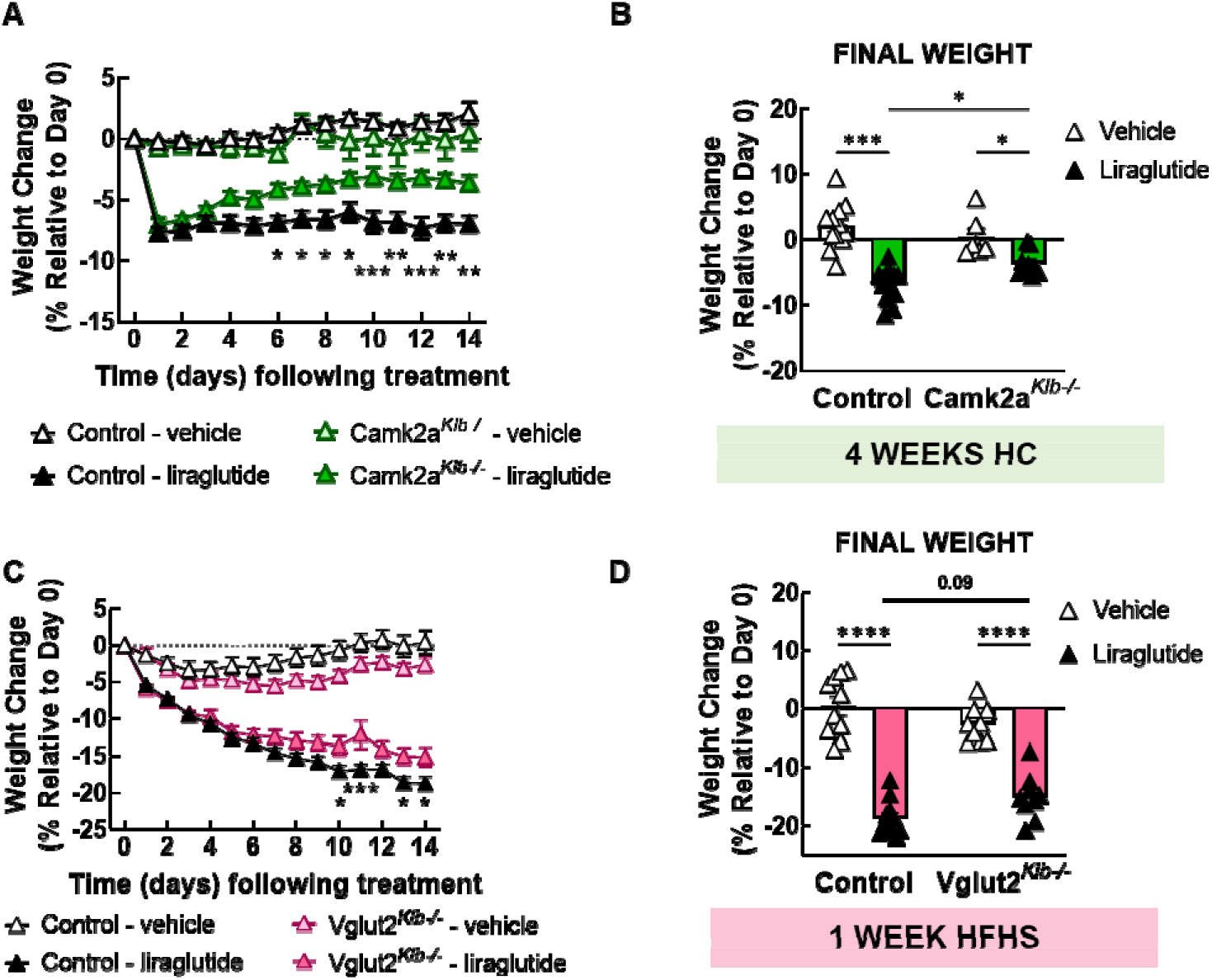
Central *Fgf21* signaling facilitates the weight-lowering effects of liraglutide. (**A-B**) Relative weight loss over time (A; Mixed-effects analysis: Interactions, F (14, 294) = 3.552, P <0.0001) and on the last day of treatment (B; Mixed-effects analysis: Interactions, F (1, 38) = 7.686, P = 0.009) in control and CAMK2A+ neuron-*Klb* knockout (CamK2a^*Klb-/-*^) mice fed a low-fat, high-carbohydrate diet for 4 weeks and then treated with vehicle or liraglutide (200 μg/kg, *b.i.d*) for 14 days (N = 9-15). (**C-D**) Relative weight loss over time (A; Mixed-effects analysis: Interactions, F (14, 252) = 5.031, P <0.0001) and on the last day of treatment (B; Mixed-effects analysis: Interactions, F (1, 35) = 7.296, P = 0.0106) in control and vGLUT2+ neuron-*Klb* knockout (Vglut2^*Klb-/-*^) mice fed a high-fat, high-sugar diet for 1 week and then treated with vehicle or liraglutide (200 μg/kg, *b.i.d*) for 14 days (N = 9-11). Data are shown as mean ± SEM, ns not significant, * P < 0.05, ** P < 0.01, *** P < 0.001, and **** P < 0.0001 for comparisons delineated by lines (B, D) and liraglutide-treated control vs. Liv^*Fgf21-/-*^ mice (A, C).

## Discussion

In this study, we identify a novel brain GLP-1R-liver FGF21 axis that mediates the weight-lowering effect of GLP-1RA in the presence of diets with high carbohydrate content. These results could provide insight into the clinically observed variability in weight loss following GLP-1R agonist treatment. While liraglutide and semaglutide can promote up to 10% and 15% weight loss, respectively, many individuals lose significantly less weight in response to these drugs^39–41^. Results from this preclinical study suggest the hypothesis that macronutrient content and FGF21-related factors (e.g., FGF21-resistant states, FGF21 polymorphisms) could influence the weight loss efficacy of GLP-1RA and thereby may help explain the significant heterogeneity in the magnitude of weight loss following GLP-1RA treatment in humans.

GLP-1RA such as exendin-4 and liraglutide have been previously shown to increase circulating levels of FGF21^30–35^. However, these studies did not address whether GLP-1RA stimulation of FGF21 production arises indirectly from GLP-1RA-induced reduction in food intake – a known stimulus of hepatic FGF21 secretion^36–38^. Here, we demonstrate that liraglutide increases FGF21 production independently of its effects on food intake. Although the liver is the primary source of circulating FGF21^38^, the GLP-1R is not expressed in hepatocytes^42–44^. This strongly suggests an indirect mechanism for GLP-1R agonist-mediated stimulation of FGF21 levels. Utilizing multiple tissue-specific knockout mouse models, we show that neuronal GLP-1R expression, specifically in glutamatergic neurons and those within the cellular target domains of *Wnt1Cre2*, is required for the stimulatory effect of liraglutide on FGF21. These findings provide an important foundation for future studies investigating how neuronal GLP-1R activation stimulates FGF21 production by the liver. One possibility is that GLP-1RA-induced liver FGF21 production is mediated by direct autonomic projections from the brain to the liver. Central GLP-1R activation has also been shown to increase sympathetic outflow to adipose depots in rodents^57–60^, which could stimulate lipolysis and subsequent release of free fatty acids, a potent stimulator of liver PPARα and FGF21 production. Another potential mechanism by which central GLP-1R activation could stimulate hepatic FGF21 production is via the hypothalamic-pituitary-adrenal (HPA) axis. It is well-established that GLP-1RA, administered centrally or peripherally, activate the HPA axis to increase circulating corticosterone levels in both rodents and humans^61–63^. Glucocorticoids (GC) can in turn induce *Fgf21* expression in the liver via hepatic GC receptor activation^64^. Future studies utilizing pharmacological and surgical methods to disrupt these pathways will be important to delineate the role of autonomic innervation and/or the HPA axis in mediating the GLP-1R-FGF21 interaction.

Our finding that liraglutide-induced reduction in body weight is attenuated in chow-fed *Liv^Fgf21-/-^* mice indicates that FGF21 is a component of the anorectic effect of GLP-1RA. Among its many metabolic actions, FGF21 plays an important role in regulating carbohydrate intake and preference. Liver FGF21 production is stimulated by high carbohydrate intake^29,65,66^, and this, in turn, acts as a feedback inhibitor of subsequent carbohydrate consumption^28,29^. Here we hypothesized that GLP-1RA-induced FGF21 acts in a similar manner and facilitates the decreased intake of carbohydrate-rich diets in response to GLP-1RA treatment. This hypothesis was supported by our finding that loss of liver *Fgf21* expression selectively attenuated liraglutide-induced weight loss in mice fed a high-carbohydrate diet. Interestingly, we also demonstrate that in control mice, liraglutide reduces body weight to a greater degree in mice fed a high-carbohydrate diet. These findings may be clinically relevant since they raise the testable hypothesis that dietary carbohydrate content can modify the effectiveness of GLP-1RA as weight loss drugs. Moreover, since several variants in the *hFGF21* gene locus have been associated with effects on carbohydrate intake and sweet preference in humans^45–50^, results from our studies propose a potential significance of these genetic variants in influencing an individual’s weight loss response to GLP-1RA.

Weight loss in response to GLP-1RA is primarily attributed to reduced food intake. Although GLP-1RA stimulate sympathetic outflow to adipose tissues in rodents^57–59^, a consistent effect on energy expenditure has not been clearly established^67^. On the other hand, pharmacological levels of FGF21 have been shown to increase energy expenditure in mice^21,25^. A previous study reported that liraglutide stimulates FGF21 production from adipose tissue-resident invariant natural killer T (iNKT) cells, and that this increased FGF21, in turn, promoted weight loss via increased energy expenditure^33^. In the present studies, we did not observe increased energy expenditure in response to liraglutide in control or Liv^*Fgf21*-/-^ mice but instead observed a decrease in energy expenditure in both groups. Interestingly, liraglutide-treated Liv^*Fgf21*-/-^ mice displayed a greater decrease in energy expenditure compared to control mice. This suggests that an increase in food intake and a greater reduction in energy expenditure additively contribute to the attenuation of the weight-lowering effect of liraglutide in Liv^*Fgf21*-/-^ mice. Future studies will address the target tissues and mechanisms mediating the effects of liraglutide-induced increases in FGF21 levels on food intake and energy expenditure.

FGF21 signals via a FGFR1c-KLB dimer. Since the tissue specificity of FGF21 actions is conferred by expression of KLB, site-specific knockouts of *Klb* are used to disrupt FGF21 signaling in different cell types and tissues^51–53^. Disruption of *Klb* in forebrain regions expressing *Camk2a* render mice unresponsive to the pharmacological effects of FGF21 on body weight^55^ and sweet preference^28^. In addition, *Klb* expression in glutamatergic neurons mediates the suppressive effect of FGF21 on carbohydrate intake^56^ and body weight^68,69^. Our finding that liraglutide-induced weight loss is also diminished in mice lacking *Klb* in *Camk2a*-expressing cells suggests that forebrain neurons are targeted by FGF21 to reduce body weight in response to liraglutide. We further demonstrate that loss of *Klb* expression in glutamatergic neurons also attenuates liraglutide-induced weight loss in mice fed a high-fat, high-sugar diet. This proposes that liraglutide-induced FGF21 promotes weight loss in the presence of high carbohydrate diets via its actions in glutamatergic neurons in forebrain regions of the central nervous system. Future studies will use region-specific *Klb* knockout models to more precisely identify the brain region(s) mediating this effect. A key target is the ventromedial hypothalamus since loss of *Klb* expression in this brain region blocks the ability of FGF21 to reduce carbohydrate intake and sucrose preference^56^. The paraventricular nucleus of the hypothalamus is another potential site of liraglutide-induced FGF21 actions as loss of *Klb* expression in this region attenuates baseline preference for sucrose even in the absence of markedly increased FGF21^56^.

The present studies identify FGF21 as a component of a novel brain-liver-brain crosstalk that plays a key role in mediating the food intake- and weight-suppressive benefits of the GLP-1RA liraglutide in the presence of high carbohydrate diets. More in-depth preclinical and clinical studies into the role of the brain GLP-1R-liver FGF21 crosstalk may shed light on the well-documented variability in response to GLP-1RA and thereby enable precision medicine tailoring of GLP-1-based therapeutics to different individuals based on their genetics and environment such as diet. Given the benefits and growing call for wider implementation of “food is medicine” interventions such as medically tailored meals (MTMs) for diet-related diseases^70,71^, more indepth understanding of how dietary composition modifies GLP-1RA efficacy would inform refinement of current and future therapeutic protocols for the use of MTMs and GLP-1-based therapeutics for chronic weight management. Moreover, our results support the novel notion that the anorectic effect of GLP-1RA is comprised of discrete and differentially regulated actions of these compounds influenced by different dietary components. Better understanding of these pathways may drive development of novel strategies such as dual and/or biased agonists^72^ to fully harness the therapeutic potential of the GLP-1 system. Lastly, while the current study focuses on food intake and body weight, GLP-1RA and FGF21 analogues share many other metabolic benefits including protection against cardiovascular diseases^73,74^, hepatosteatosis^75,76^, neurodegenerative diseases^77–79^, and suppression of alcohol consumption^28,80–82^. Future studies examining the potential role of the GLP-1R-FGF21 axis in these therapeutic areas could inform the development of novel pharmacologic strategies for the treatment of these conditions.

## Methods

### Animal models and husbandry

C57BL/6J male mice (Cat. No. 664, The Jackson Laboratory, Inc.) were used in FGF21 measurement studies. Vglut2^*Glp1r-/-*^ and Wnt1^*Glp1r-/-*^ mice were generated by crossing *vGlut2-Cre* or *Wnt1-Cre* mice, respectively, with *floxed-Glp1r* mice as described previously^13,14^. β cell^*Glp1r-/-*^ mice were generated by crossing MIP-CreERT with *floxed-Glp1r* mice as previously described^83^. Liv^*Ppar-/-*^ mice were generated by crossing *Alb*-Cre mice with *floxed-Pppara* mice (a generous gift from Dr. Dan Kelly, University of Pennsylvania). *Liv^fgf21-/-^* mice were generated by crossing *Alb*-Cre mice with *floxed-Fgf21* mice (Cat. No. 22361, The Jackson Laboratory, Inc.). Camk2a^*Klb-/-*^ and Vglut2^*Klb-/-*^ mice were generated by crossing *Camk2a-Cre* and *vGlut2-Cre* mice, respectively with *floxed-Klb* mice (*Camk2a*-Cre and *floxed-Klb* mice were a generous gift from Dr. Steven Kliewer, University of Texas Southwestern, and were provided by Dr. Christopher Morrison, Pennington Biomedical Research Center), as previously described^55,84^. All mice were housed on a standard 12 h/12 h light/dark cycle. They had *ad libitum* access to distilled water and were maintained on a chow diet (57.9% calories provided by carbohydrates, 28.7% protein, 13.4% fat; 3.36 kcal/g; 5L0D, LabDiet, St. Louis, MO) from the time of weaning unless specified otherwise.

### Acute liraglutide administration

Weight-matched C57BL/6J male mice, Vglut2^*Glp1r-/-*^, β cell^*Glp1r-/-*^, Liv^*Ppar-/-*^, and Wnt1^*Glp1r-/-*^ mice and respective background-matched control mice were fasted for 4 h at the start of the light cycle and randomly assigned to receive either vehicle (0.9% saline) or liraglutide (400 μg/kg body weight) subcutaneously. Body weight, blood glucose, and tail blood were collected at 0 and 7 h following treatment.

### Pair-feeding studies

Weight-matched, adult C57BL/6J male mice were randomly assigned to receive either vehicle (0.9% saline) or liraglutide (200 μg/kg body weight) subcutaneously twice a day while having *ad libitum* access to food or while being pair-fed to weight-match liraglutide-treated mice for 48 h. Food intake and body weight were monitored throughout the study period. Tail blood was collected at 0 and 48 h following treatment.

### Metabolic chamber experiments with chronic liraglutide administration

Two cohorts (16 mice each) of 15-17-week-old weight-matched control and *Liv^Fgf21-/-^* male mice fed a chow diet were individually housed for 5-7 days before being placed in a Promethion metabolic system (Sable Systems, Inc.). Following a 5-7 day acclimation period, mice were randomly assigned to receive vehicle (0.9% saline) or liraglutide (200 μg/kq body weight) twice a day for 12 days. Food intake, energy expenditure, and other metabolic parameters (water intake, respiratory exchange ratio, activity) were recorded continuously during the study period. Body weight was measured manually every day. Body composition measurements were obtained by NMR (Minispec 235 LF90II-TD NMR Analyzer, Bruker) at the start and end of the treatment period.

### Low and high carbohydrate diet experiments

10-12-week-old control and *Liv^Fgf21-/-^* male mice were placed on a low-fat, low-carbohydrate or low-fat, high-carbohydrate diet (D08091802 or D12450J, Research Diets, Inc., respectively) for 4 weeks. Separate cohorts of 10-12-week-old and 14-15-week-old control and *Liv^Fgf21-/-^* male mice were placed on a high-fat, high-sugar diet (D12451, Research Diets, Inc.) for 4 weeks or 1 week, respectively. Mice were then randomly assigned to receive vehicle (0.9% saline) or liraglutide (200 μg/kg body weight) subcutaneously twice a day for 14 days while being maintained on their respective diet. Body weight was measured daily. Body composition was measured at the start and end of the treatment period.

### Central Klb knockout studies

10-12-week-old control and Camk2a^*Klb-/-*^ male mice were placed on a low-fat, high-carbohydrate diet (D12450J, Research Diets, Inc.) for 4 weeks. 14-15-week-old control and Vglut2^*Klb-/-*^ were placed on a high-fat, high-sugar diet (D12451, Research Diets, Inc.) for 1 week. Mice were then randomly assigned to receive vehicle (0.9% saline) or liraglutide (200 μg/kg body weight) subcutaneously twice a day for 14 days while being maintained on their respective diet. Body weight was measured daily. Body composition was measured at the start and end of the treatment period.

### FGF21 measurements

Tail blood was collected in EDTA-coated microvette tubes (16.444.100, Sarstedt Inc.) and centrifuged at 4,000 x g at 4°C for 20 minutes. Plasma was collected and stored at −20°C until analysis. Plasma FGF21 was measured using a commercially available mouse FGF21 ELISA kit (ab212160, Abcam).

### RNA isolation and qPCR

Total RNA was extracted from tissues using reagents and following instructions from the DirectZol RNA Mini-prep (R2051, Zymogen). cDNA was synthesized by reverse transcription using the iScript cDNA Synthesis Kit (1708891, Bio-Rad). Real time PCR reactions were performed using TaqMan Real-Time PCR Assays (Mm00840165_g1, Thermo Fisher Scientific) and TaqMan Fast Advanced Master Mix (4444556, Applied Biosystems).

### Statistics

Data were analyzed using GraphPad Prism 9 Software (GraphPad Software, Inc., La Jolla, CA). One-way ANOVA or mixed-effects analysis followed by Holm-Sidak’s multiple comparisons was used when appropriate and indicated in the figure legends. P < 0.05 was considered statistically significant. Values represent mean ± SEM. Energy expenditure (EE) data were analyzed using the EE analysis of covariance (ANCOVA) analysis provided by the NIDDK Mouse Metabolic Phenotyping Centers (MMPC, www.mmpc.org) using their Energy Expenditure Analysis page (http://www.mmpc.org/shared/regression.aspx) and supported by grants DK076169 and DK115255.

### Study approval

All animal studies were approved by the Institutional Animal Care and Use Committees at Vanderbilt University, University of Michigan, Duke University, University of Iowa, and the Toronto Center for Phenogenomics, Mt. Sinai Hospital.

## Author contributions

TDVL and JEA designed the experiments and wrote the manuscript. TDVL, PF, ABW, BJE, NB-K, LLB, JK, JLB, MBB, MBP, JPR, AIS and KGN performed the experiments and/or analyzed the data. TDVL, JEC, LLB, DJD, NB-K, RJS, MJP and JEA edited the manuscript. All authors reviewed the final version.

## Acknowledgements

We thank Dr. Steven Kliewer and Dr. Christopher Morrison for the generous donation of the Camk2a^*Klb-/-*^ mice and Dr. Dan Kelly for the generous donation of the *floxed-Ppara* mice. We also thank Dr. Louise Lantier and Merrygay James of the Vanderbilt Mouse Metabolic Phenotyping Center (VMMPC) for assisting with the metabolic chamber experiments. The VMMPC is supported by the National Institutes of Health (5U2CDK059637). Research reported in this publication was supported by National Institutes of Health (NIH) award T32GM007347 (TDVL); NIH award R01DK106104 (MJP); NIH awards R01DK123075, R01DK125353, and R01DK046492 (JEC); CIHR grant 154321 (DJD); and Department of Veterans Affairs IK2BX005715 (NB-K). DJD is also supported by a Banting and Best Diabetes Centre chair in Incretin Biology and the Sinai Health-Novo Nordisk Foundation Fund in regulatory peptides. JEC is also supported by the Helmsley Charitable Trust Foundation.

---

**Supplemental Figure 1.**
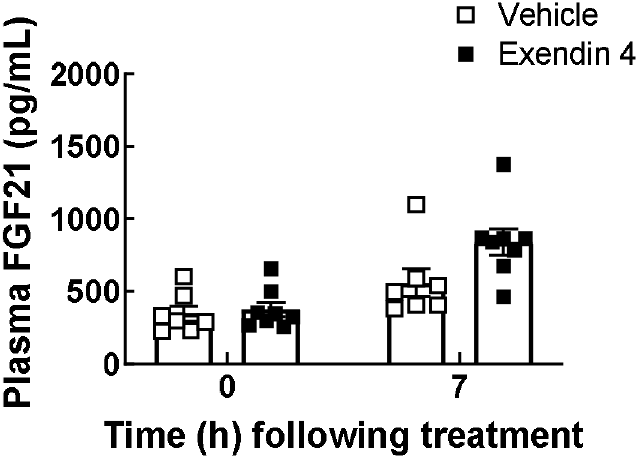
Exendin-4 stimulates FGF21 in fasted mice. Plasma FGF21 levels in mice treated with Exendin-4 (100 μg/kg) following the study protocol shown in **Figure 1A**. (Mixed-effects analysis: Interaction, F (1, 26) = 2.93, p = 0.0987; Treatment Effect, F (1,26) = 4.30, P = 0.0481; N = 8).

**Supplemental Figure 2.**
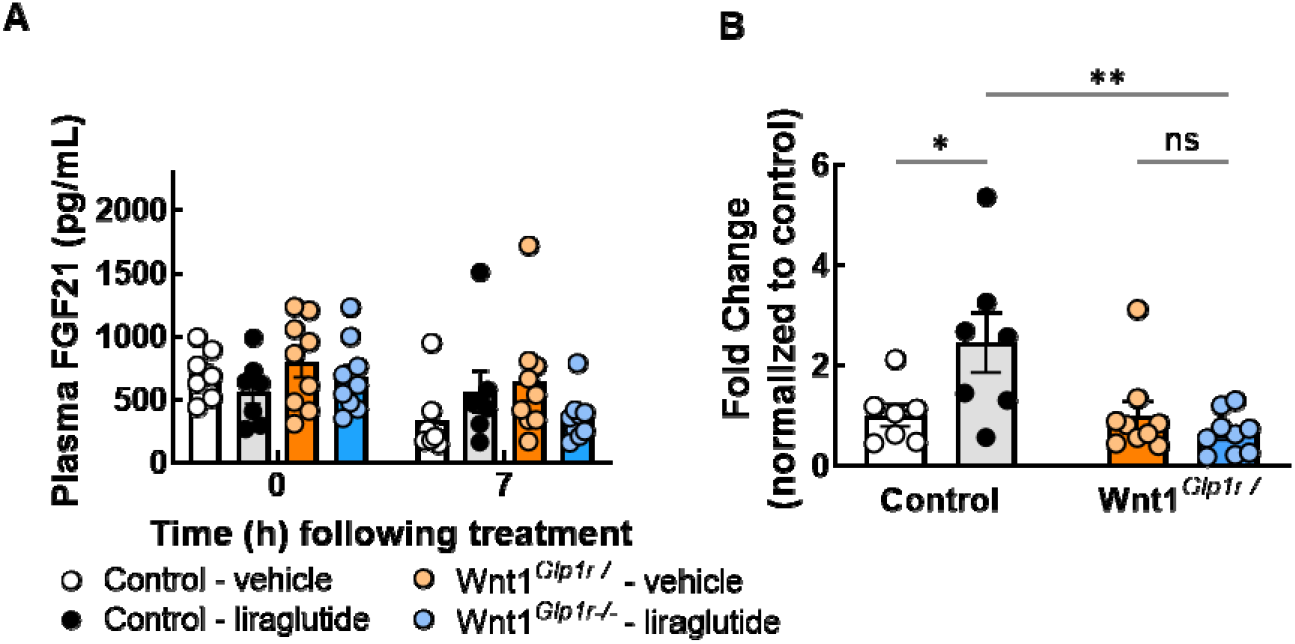
The GLP-1R in Wnt1+ cells is required for the increase in plasma FGF21 levels following liraglutide treatment. Absolute (**A**) and relative change (**B**) in FGF21 levels at 0 and 7 h post treatment with vehicle or liraglutide (400 μg/kg) in fasted control mice and mice lacking the GLP-1R in Wnt1+ neurons (Wnt1^*Glp1r-/-*^) (A, Mixed-effects analysis: Interaction, F (1, 28) = 4.131, P = 0.0517; Genotype Effect, F (1, 28) = 0.6444, P = 0.4289; B, Mixed-effects analysis: Interaction, F (1, 28) = 7.458, P = 0.0108; N = 7-9). Data are shown as mean ± SEM, ns not significant, * P < 0.05, ** P < 0.01.

**Supplemental Figure 3.**
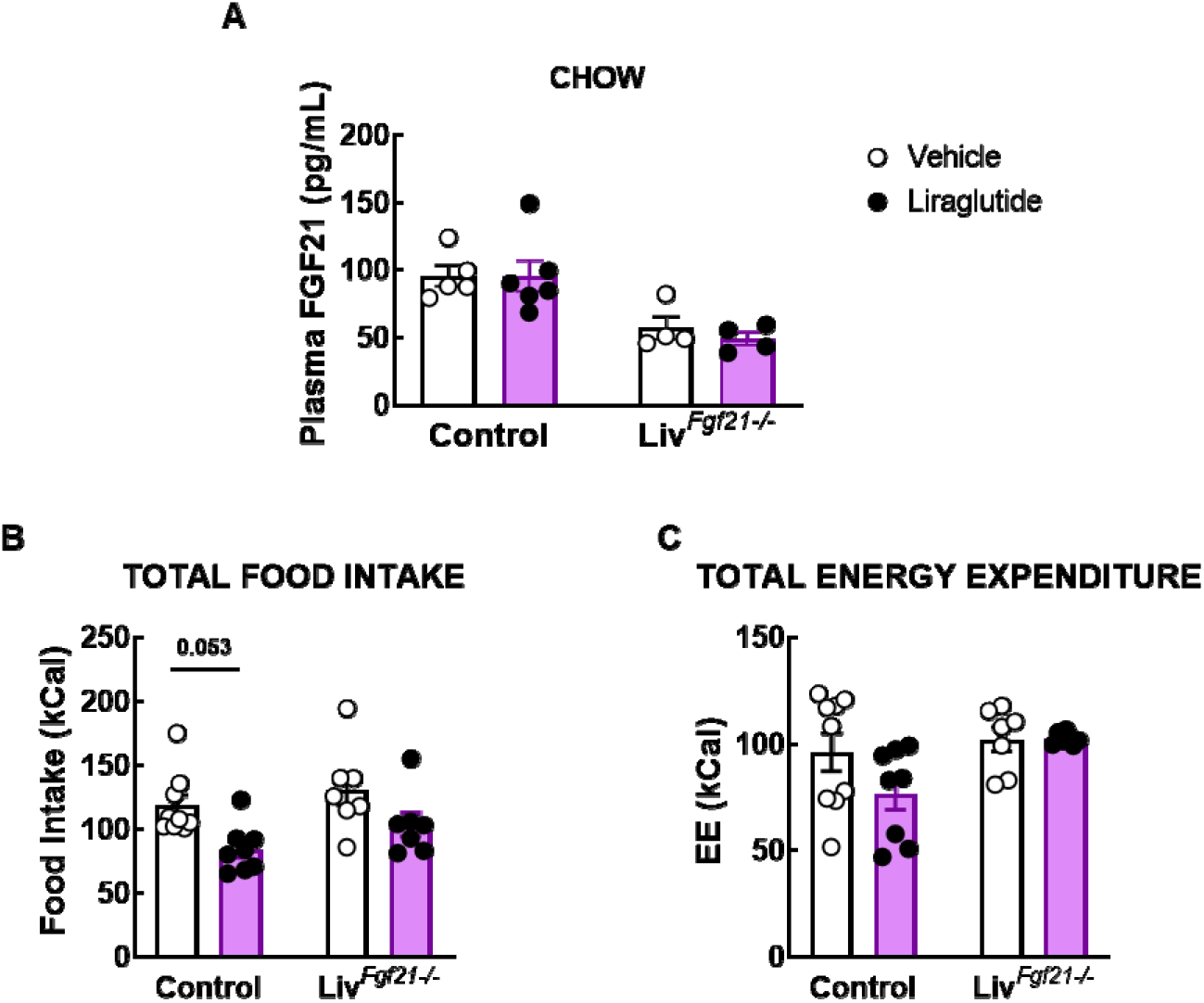
Loss of liver FGF21 expression attenuates the anorectic effect of liraglutide. (**A**) Plasma FGF21 levels in control and Liv^*Fgf21-/-*^ mice following treatment with vehicle or liraglutide (200 μg/kg, *b.i.d*) for 12 days (Mixed-effects analysis: Interaction, F (1, 15) = 31.45, P < 0.0001; Genotype Effect, F (1, 15) = 92.74, P < 0.0001; N = 5-6). (**B**) Cumulative food intake (Mixed-effects analysis: Interaction, F (1, 27) = 0.1371, P = 0.7140; Genotype Effect, F (1, 27) = 2.691, P = 0.1125; N = 7-9) and (**C**) cumulative EE (Mixed-effects analysis: Interaction, F (1, 27) = 2.086, P = 0.1601; Genotype Effect, F (1, 27) = 5.068, P=0.0327; N = 7-9) in control and *Liv^Fgf21-/-^* mice following treatment with vehicle or liraglutide (200 μg/kg, *b.i.d*) for 12 days in metabolic cages. Data are shown as mean ± SEM.

**Supplemental Figure 4.**
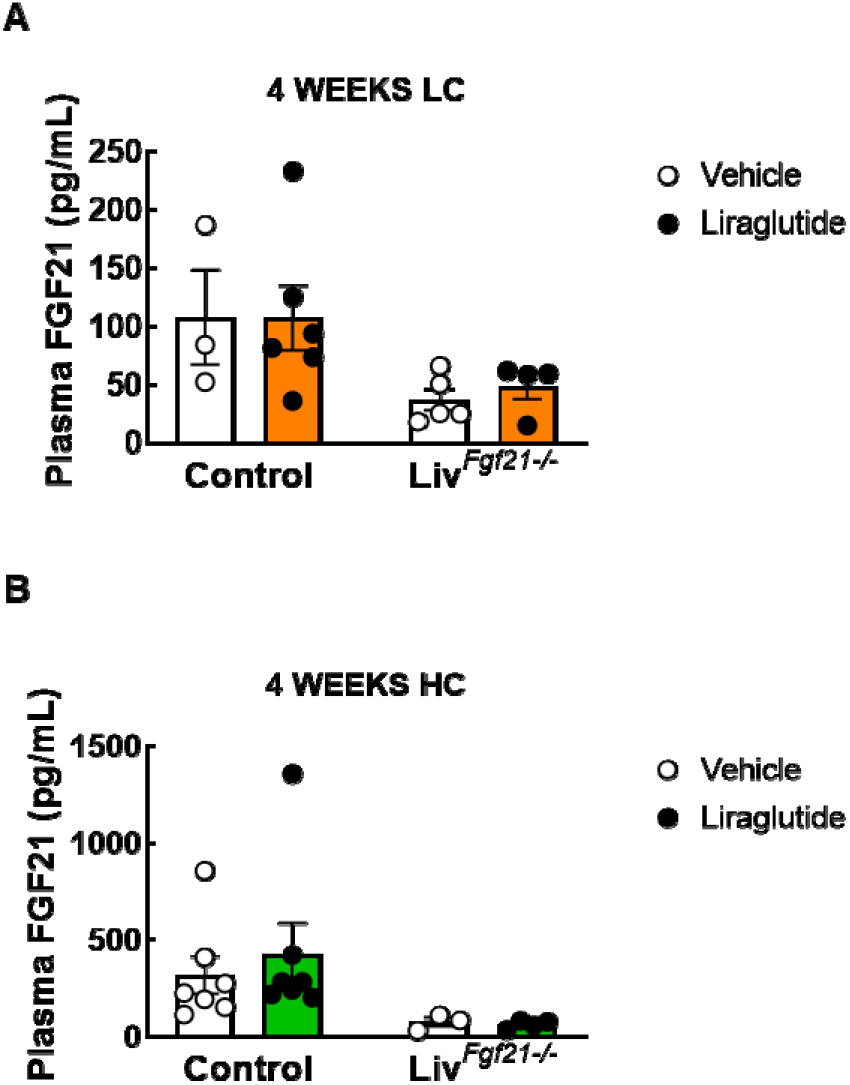
Plasma FGF21 in LC and HC-fed mice. Plasma FGF21 levels in (**A**) LC- and (**B**) HC-fed control and Liv^*Fgf21-/-*^ mice following treatment with vehicle or liraglutide (200 μg/kg, *b.i.d*) for 12 days (A; Mixed-effects analysis: Interaction, F (1, 14) = 3.111, P = 0.0996; Genotype Effect, F (1, 14) = 21.25, P=0.0004; N = 3-6 and B; Mixed-effects analysis: Interaction, F (1, 17) = 1.353, P = 0.2609; Genotype Effect, F (1, 17) = 7.268, P=0.0153; N = 4-7.

## References

1. Fryar CD, Carroll MD, Afful J. Prevalence of Overweight, Obesity, and Extreme Obesity Among Adults Aged 20 and Over: United States, 1960-1962 Through 2017-2018. NCHS Health E-Stats. Published online 2020.

2. Stierman B, Afful J, Carroll MD, et al. National Health and Nutrition Examination Survey 2017-March 2020 Prepandemic Data Files Development of Files and Prevalence Estimates for Selected Health Outcomes. Natl Health Stat Rep. 2021;2021(158). doi:10.15620/CDC:106273

3. Astrup A, Rössner S, Van Gaal L, et al. Effects of liraglutide in the treatment of obesity: a randomised, double-blind, placebo-controlled study. The Lancet. 2009;374(9701):1606–1616. doi:10.1016/S0140-6736(09)61375-1

4. Vilsbøll T, Christensen M, Junker AE, Knop FK, Gluud LL. Effects of glucagon-like peptide-1 receptor agonists on weight loss: systematic review and meta-analyses of randomised controlled trials. BMJ. 2012;344(7841). doi:10.1136/BMJ.D7771

5. Pi-Sunyer X, Astrup A, Fujioka K, et al. A Randomized, Controlled Trial of 3.0 mg of Liraglutide in Weight Management. N Engl J Med. 2015;373(1):11–22. doi:10.1056/NEJMOA1411892/SUPPL_FILE/NEJMOA1411892_DISCLOSURES.PDF

6. Davies M, Færch L, Jeppesen OK, et al. Semaglutide 2·4 mg once a week in adults with overweight or obesity, and type 2 diabetes (STEP 2): a randomised, double-blind, double-dummy, placebo-controlled, phase 3 trial. The Lancet. 2021;397(10278):971–984. doi:10.1016/S0140-6736(21)00213-0

7. Wadden TA, Bailey TS, Billings LK, et al. Effect of Subcutaneous Semaglutide vs Placebo as an Adjunct to Intensive Behavioral Therapy on Body Weight in Adults With Overweight or Obesity: The STEP 3 Randomized Clinical Trial. JAMA. 2021;325(14):1403–1413. doi:10.1001/JAMA.2021.1831

8. Wilding JPH, Batterham RL, Calanna S, et al. Once-Weekly Semaglutide in Adults with Overweight or Obesity. N Engl J Med. 2021;384(11):989–1002. doi:10.1056/NEJMOA2032183/SUPPL_FILE/NEJMOA2032183_DATA-SHARING.PDF

9. Vosoughi K, Atieh J, Khanna L, et al. Association of Glucagon-like Peptide 1 Analogs and Agonists Administered for Obesity with Weight Loss and Adverse Events: A Systematic Review and Network Meta-analysis. EClinicalMedicine. 2021;42:101213.

10. Secher A, Jelsing J, Baquero AF, et al. The arcuate nucleus mediates GLP-1 receptor agonist liraglutide-dependent weight loss. J Clin Invest. 2014;124(10):4473–4488. doi:10.1172/JCI75276

11. Sisley S, Gutierrez-Aguilar R, Scott M, D’Alessio DA, Sandoval DA, Seeley RJ. Neuronal GLP1R mediates liraglutide’s anorectic but not glucose-lowering effect. J Clin Invest.2014;124(6):2456–2463. doi:10.1172/JCI72434

12. Burmeister MA, Ayala JE, Smouse H, et al. The hypothalamic glucagon-like peptide 1 receptor is sufficient but not necessary for the regulation of energy balance and glucose homeostasis in mice. Diabetes. 2017;66(2):372–384. doi:10.2337/db16-1102

13. Adams JM, Pei H, Sandoval DA, et al. Liraglutide modulates appetite and body weight through glucagon-like peptide 1 receptor-expressing glutamatergic neurons. In: Diabetes.Vol 67. American Diabetes Association Inc.; 2018:1538–1548. doi:10.2337/db17-1385

14. Varin EM, Mulvihill EE, Baggio LL, et al. Distinct Neural Sites of GLP-1R Expression Mediate Physiological versus Pharmacological Control of Incretin Action. Cell Rep.2019;27(11):3371–3384.e3. doi:10.1016/J.CELREP.2019.05.055

15. Potthoff MJ, Kliewer SA, Mangelsdorf DJ. Endocrine fibroblast growth factors 15/19 and 21: from feast to famine. Genes Dev. 2012;26(4):312–324. doi:10.1101/GAD.184788.111

16. Fisher FM, Maratos-Flier E. Understanding the Physiology of FGF21. Annu Rev Physiol. 2016;78:223–241. doi:10.1146/ANNUREV-PHYSIOL-021115-105339

17. Kharitonenkov A, DiMarchi R. Fibroblast growth factor 21 night watch: advances and uncertainties in the field. J Intern Med. 2017;281(3):233–246.

18. BonDurant LD, Potthoff MJ. Fibroblast Growth Factor 21: A Versatile Regulator of Metabolic Homeostasis. Annu Rev Nutr. 2018;38(1):annurev-nutr-071816-064800. doi:10.1146/annurev-nutr-071816

19. Kliewer SA, Mangelsdorf DJ. A Dozen Years of Discovery: Insights into the Physiology and Pharmacology of FGF21. Cell Metab. 2019;29(2):246–253. doi:10.1016/J.CMET.2019.01.004

20. Hill CM, Qualls-Creekmore E, Berthoud HR, et al. FGF21 and the Physiological Regulation of Macronutrient Preference. Endocrinology. 2020;161(3). doi:10.1210/ENDOCR/BQAA019

21. Xu J, Lloyd DJ, Hale C, et al. Fibroblast Growth Factor 21 Reverses Hepatic Steatosis, Increases Energy Expenditure, and Improves Insulin Sensitivity in Diet-Induced Obese Mice. Diabetes. 2009;58(1):250–259. doi:10.2337/db08-0392

22. Kharitonenkov A, Shiyanova TL, Koester A, et al. FGF-21 as a novel metabolic regulator. J Clin Invest. 2005;115(6):1627–1635. doi:10.1172/JCI23606

23. Davies MJ, Bergenstal R, Bode B, et al. Efficacy of Liraglutide for Weight Loss Among Patients With Type 2 Diabetes: The SCALE Diabetes Randomized Clinical Trial. JAMA. 2015;314(7):687–699. doi:10.1001/JAMA.2015.9676

24. Kaufman A, Abuqayyas L, Denney WS, Tillman EJ, Rolph T. AKR-001, an Fc-FGF21 Analog, Showed Sustained Pharmacodynamic Effects on Insulin Sensitivity and Lipid Metabolism in Type 2 Diabetes Patients. Cell Rep Med. 2020;1(4):100057. doi:10.1016/J.XCRM.2020.100057

25. Coskun T, Bina HA, Schneider MA, et al. Fibroblast Growth Factor 21 Corrects Obesity in Mice. Endocrinology. 2008;149(12):6018–6027. doi:10.1210/en.2008-0816

26. Gaich G, Chien JY, Fu H, et al. The Effects of LY2405319, an FGF21 Analog, in Obese Human Subjects with Type 2 Diabetes. Cell Metab. 2013;18(3):333–340. doi:10.1016/j.cmet.2013.08.005

27. Talukdar S, Zhou Y, Li D, et al. A Long-Acting FGF21 Molecule, PF-05231023, Decreases Body Weight and Improves Lipid Profile in Non-human Primates and Type 2 Diabetic Subjects. Cell Metab. 2016;23(3):427–440. doi:10.1016/J.CMET.2016.02.001

28. Talukdar S, Owen BM, Mangelsdorf DJ, et al. FGF21 Regulates Sweet and Alcohol Preference Cell Metabolism FGF21 Regulates Sweet and Alcohol Preference. Cell Metab.2016;23:344–349. doi:10.1016/j.cmet.2015.12.008

29. Von Holstein-Rathlou S, Bondurant LD, Peltekian L, et al. FGF21 mediates endocrine control of simple sugar intake and sweet taste preference by the liver. Cell Metab.2016;23(2):335–343. doi:10.1016/j.cmet.2015.12.003

30. Yang M, Zhang L, Wang C, et al. Liraglutide Increases FGF-21 Activity and Insulin Sensitivity in High Fat Diet and Adiponectin Knockdown Induced Insulin Resistance. Bhattacharya S, ed. PLoS ONE. 2012;7(11):e48392. doi:10.1371/journal.pone.0048392

31. Nonogaki K, Hazama M, Satoh N. Liraglutide Suppresses Obesity and Hyperglycemia Associated with Increases in Hepatic Fibroblast Growth Factor 21 Production in KKA y Mice. BioMed Res Int. 2014;2014:1–8. doi:10.1155/2014/751930

32. Lee J, Hong SW, Park SE, et al. Exendin-4 regulates lipid metabolism and fibroblast growth factor 21 in hepatic steatosis. Metabolism. 2014;63(8):1041–1048. doi:10.1016/J.METABOL.2014.04.011

33. Lynch L, Hogan AE, Duquette D, et al. iNKT Cells Induce FGF21 for Thermogenesis and Are Required for Maximal Weight Loss in GLP1 Therapy. Cell Metab. 2016;24(3):510–519. doi:10.1016/J.CMET.2016.08.003

34. Liu J, Yang K, Yang J, et al. Liver-derived fibroblast growth factor 21 mediates effects of glucagon-like peptide-1 in attenuating hepatic glucose output. EBioMedicine. 2019;41:73–84. doi:10.1016/J.EBIOM.2019.02.037

35. Liu D, Pang J, Shao W, et al. Hepatic fibroblast growth factor 21 is involved in mediating functions of liraglutide in mice with dietary challenge. Hepatology. 2021;74(4):2154–2169. doi:10.1002/hep.31856

36. Badman MK, Pissios P, Kennedy AR, Koukos G, Flier JS, Maratos-Flier E. Hepatic Fibroblast Growth Factor 21 Is Regulated by PPARα and Is a Key Mediator of Hepatic Lipid Metabolism in Ketotic States. Cell Metab. 2007;5(6):426–437. doi:10.1016/j.cmet.2007.05.002

37. Inagaki T, Dutchak P, Zhao G, et al. Endocrine Regulation of the Fasting Response by PPARα-Mediated Induction of Fibroblast Growth Factor 21. Cell Metab. 2007;5(6):415–425. doi:10.1016/j.cmet.2007.05.003

38. Markan KR, Naber MC, Ameka MK, et al. Circulating FGF21 is liver derived and enhances glucose uptake during refeeding and overfeeding. Diabetes. 2014;63(12):4057–4063. doi:10.2337/db14-0595

39. O’Neil PM, Birkenfeld AL, McGowan B, et al. Efficacy and safety of semaglutide compared with liraglutide and placebo for weight loss in patients with obesity: a randomised, doubleblind, placebo and active controlled, dose-ranging, phase 2 trial. Lancet Lond Engl.2018;392(10148):637–649. doi:10.1016/S0140-6736(18)31773-2

40. Capehorn MS, Catarig AM, Furberg JK, et al. Efficacy and safety of once-weekly semaglutide 1.0mg vs once-daily liraglutide 1.2mg as add-on to 1-3 oral antidiabetic drugs in subjects with type 2 diabetes (SUSTAIN 10). Diabetes Metab. 2020;46(2):100–109. doi:10.1016/J.DIABET.2019.101117

41. Garvey WT, Batterham RL, Bhatta M, et al. Two-year effects of semaglutide in adults with overweight or obesity: the STEP 5 trial. Nat Med. 2022;28(10):2083–2091. doi:10.1038/S41591-022-02026-4

42. Panjwani N, Mulvihill EE, Longuet C, et al. GLP-1 Receptor Activation Indirectly Reduces Hepatic Lipid Accumulation But Does Not Attenuate Development of Atherosclerosis in Diabetic Male ApoE-/- Mice. Endocrinology. 2013;154(1):127–139. doi:10.1210/en.2012-1937

43. McLean BA, Wong CK, Kaur KD, Seeley RJ, Drucker DJ. Differential importance of endothelial and hematopoietic cell GLP-1Rs for cardiometabolic versus hepatic actions of semaglutide. JCI Insight. 2021;6(22). doi:10.1172/JCI.INSIGHT.153732

44. McLean BA, Wong CK, Campbell JE, Hodson DJ, Trapp S, Drucker DJ. Revisiting the Complexity of GLP-1 Action from Sites of Synthesis to Receptor Activation. Endocr Rev.2021;42(2):101–132. doi:10.1210/endrev/bnaa032

45. Chu AY, Workalemahu T, Paynter NP, et al. Novel locus including FGF21 is associated with dietary macronutrient intake. Hum Mol Genet. 2013;22(9):1895–1902. doi:10.1093/HMG/DDT032

46. Tanaka T, Ngwa JS, Van Rooij FJA, et al. Genome-wide meta-analysis of observational studies shows common genetic variants associated with macronutrient intake. Am J Clin Nutr. 2013;97(6):1395–1402. doi:10.3945/AJCN.112.052183

47. Frayling TM, Beaumont RN, Jones SE, et al. A Common Allele in FGF21 Associated with Sugar Intake Is Associated with Body Shape, Lower Total Body-Fat Percentage, and Higher Blood Pressure. Cell Rep. 2018;23(2):327–336. doi:10.1016/J.CELREP.2018.03.070

48. Merino J, Dashti HS, Li SX, et al. Genome-wide meta-analysis of macronutrient intake of 91,114 European ancestry participants from the cohorts for heart and aging research in genomic epidemiology consortium. Mol Psychiatry. 2019;24(12):1920–1932. doi:10.1038/S41380-018-0079-4

49. Meddens SFW, de Vlaming R, Bowers P, et al. Genomic analysis of diet composition finds novel loci and associations with health and lifestyle. Mol Psychiatry. 2020;26(6):2056–2069. doi:10.1038/s41380-020-0697-5

50. Janzi S, González-Padilla E, Sonestedt E, et al. Single Nucleotide Polymorphisms in Close Proximity to the Fibroblast Growth Factor 21 (FGF21) Gene Found to Be Associated with Sugar Intake in a Swedish Population. Nutrients. 2021;13(11). doi:10.3390/NU13113954

51. Ogawa Y, Kurosu H, Yamamoto M, et al. βKlotho is required for metabolic activity of fibroblast growth factor 21. Proc Natl Acad Sci. 2007;104(18):7432–7437. doi:10.1073/PNAS.0701600104

52. Adams AC, Cheng CC, Coskun T, Kharitonenkov A. FGF21 Requires βklotho to Act In Vivo. PLOS ONE. 2012;7(11):e49977. doi:10.1371/JOURNAL.PONE.0049977

53. Ding X, Boney-Montoya J, Owen BM, et al. βKslotho is required for fibroblast growth factor 21 effects on growth and metabolism. Cell Metab. 2012;16(3):387–393. doi:10.1016/j.cmet.2012.08.002

54. Tacer KF, Bookout AL, Ding X, et al. Research Resource: Comprehensive Expression Atlas of the Fibroblast Growth Factor System in Adult Mouse. Mol Endocrinol. 2010;24(10):2050–2064. doi:10.1210/ME.2010-0142

55. Owen BM, Ding X, Morgan DA, et al. FGF21 acts centrally to induce sympathetic nerve activity, energy expenditure, and weight loss. Cell Metab. 2014;20(4):670–677. doi:10.1016/j.cmet.2014.07.012

56. Jensen-Cody SO, Flippo KH, Claflin KE, et al. FGF21 Signals to Glutamatergic Neurons in the Ventromedial Hypothalamus to Suppress Carbohydrate Intake. Cell Metab.2020;32(2):273–286.e6. doi:10.1016/j.cmet.2020.06.008

57. Lockie SH, Heppner KM, Chaudhary N, et al. Direct Control of Brown Adipose Tissue Thermogenesis by Central Nervous System Glucagon-Like Peptide-1 Receptor Signaling. Diabetes. 2012;61(11):2753–2762. doi:10.2337/DB11-1556

58. Beiroa D, Imbernon M, Gallego R, et al. GLP-1 Agonism Stimulates Brown Adipose Tissue Thermogenesis and Browning Through Hypothalamic AMPK. Diabetes.2014;63(10):3346–3358. doi:10.2337/DB14-0302

59. Kooijman S, Wang Y, Parlevliet ET, et al. Central GLP-1 receptor signalling accelerates plasma clearance of triacylglycerol and glucose by activating brown adipose tissue in mice. Diabetologia. 2015;58(11). Accessed February 24, 2019. /pmc/articles/PMC4589565/

60. Lee SJ, Sanchez-Watts G, Krieger JP, et al. Loss of dorsomedial hypothalamic GLP-1 signaling reduces BAT thermogenesis and increases adiposity. Mol Metab. 2018;11:33–46. doi:10.1016/J.MOLMET.2018.03.008

61. Larsen PJ, Tang-Christensen M, Jessop DS. Central Administration of Glucagon-Like Peptide-1 Activates Hypothalamic Neuroendocrine Neurons in the Rat. Endocrinology. 1997;138(10): 4445–4455. doi:10.1210/endo.138.10.5270

62. Kinzig KP, D’Alessio DA, Herman JP, et al. CNS glucagon-like peptide-1 receptors mediate endocrine and anxiety responses to interoceptive and psychogenic stressors. J Neurosci Off J Soc Neurosci. 2003;23(15):6163–6170. doi:10.1523/JNEUROSCI.23-15-06163.2003

63. Gil-Lozano M, Pérez-Tilve D, Alvarez-Crespo M, et al. GLP-1(7-36)-amide and Exendin-4 Stimulate the HPA Axis in Rodents and Humans. Endocrinology. 2010;151(6):2629–2640. doi:10.1210/en.2009-0915

64. Patel R, Bookout AL, Magomedova L, et al. Glucocorticoids Regulate the Metabolic Hormone FGF21 in a Feed-Forward Loop. Mol Endocrinol. 2015;29(2):213–223. doi:10.1210/me.2014-1259

65. Fisher ffolliott M, Kim MS, Doridot L, et al. A critical role for ChREBP-mediated FGF21 secretion in hepatic fructose metabolism. Mol Metab. 2017;6(1):14–21.

66. Lundsgaard AM, Fritzen AM, Sjøberg KA, et al. Circulating FGF21 in humans is potently induced by short term overfeeding of carbohydrates. Mol Metab. 2017;6(1):22–29. doi:10.1016/J.MOLMET.2016.11.001

67. Drucker DJ. Mechanisms of Action and Therapeutic Application of Glucagon-like Peptide-1. Vol 27. Cell Press; 2018:740–756. Accessed February 5, 2019. https://www.sciencedirect.com/science/article/pii/S1550413118301797?via%3Dihub

68. Flippo KH, Jensen-Cody SO, Claflin KE, Potthoff MJ. FGF21 signaling in glutamatergic neurons is required for weight loss associated with dietary protein dilution. Sci Rep.2020;10(1):19521. doi:10.1038/s41598-020-76593-2

69. Claflin KE, Sullivan AI, Naber MC, et al. Pharmacological FGF21 signals to glutamatergic neurons to enhance leptin action and lower body weight during obesity. Mol Metab.2022;64:101564. doi:10.1016/j.molmet.2022.101564

70. Downer S, Berkowitz SA, Berkowitz SA, Harlan TS, Olstad DL, Mozaffarian D. Food is medicine: actions to integrate food and nutrition into healthcare. BMJ. 2020;369. doi:10.1136/BMJ.M2482

71. Hager K, Cudhea FP, Wong JB, et al. Association of National Expansion of Insurance Coverage of Medically Tailored Meals With Estimated Hospitalizations and Health Care Expenditures in the US. JAMA Netw Open. 2022;5(10):e2236898-e2236898. doi:10.1001/JAMANETWORKOPEN.2022.36898

72. Baggio LL, Drucker DJ. Glucagon-like peptide-1 receptor co-agonists for treating metabolic disease. Mol Metab. 2021;46:101090. doi:10.1016/J.MOLMET.2020.101090

73. Bethel MA, Patel RA, Merrill P, et al. Cardiovascular outcomes with glucagon-like peptide-1 receptor agonists in patients with type 2 diabetes: a meta-analysis. Lancet Diabetes Endocrinol. 2018;6(2):105–113. doi:10.1016/S2213-8587(17)30412-6

74. Pan X, Shao Y, Wu F, et al. FGF21 Prevents Angiotensin II-Induced Hypertension and Vascular Dysfunction by Activation of ACE2/Angiotensin-(1-7) Axis in Mice. Cell Metab.2018;27(6):1323–1337.e5. doi:10.1016/J.CMET.2018.04.002

75. Armstrong MJ, Gaunt P, Aithal GP, et al. Liraglutide safety and efficacy in patients with non-alcoholic steatohepatitis (LEAN): a multicentre, double-blind, randomised, placebo-controlled phase 2 study. The Lancet. 2016;387(10019):679–690. doi:10.1016/S0140-6736(15)00803-X

76. Sanyal A, Charles ED, Neuschwander-Tetri BA, et al. Pegbelfermin (BMS-986036), a PEGylated fibroblast growth factor 21 analogue, in patients with non-alcoholic steatohepatitis: a randomised, double-blind, placebo-controlled, phase 2a trial. The Lancet.2018;392(10165):2705–2717. doi:10.1016/S0140-6736(18)31785-9

77. Athauda D, Maclagan K, Skene SS, et al. Exenatide once weekly versus placebo in Parkinson’s disease: a randomised, double-blind, placebo-controlled trial. The Lancet.2017;390(10103):1664–1675. doi:10.1016/S0140-6736(17)31585-4

78. Kuroda M, Muramatsu R, Maedera N, et al. Peripherally derived FGF21 promotes remyelination in the central nervous system. J Clin Invest. 2017;127(9):3496–3509. doi:10.1172/JCI94337

79. Jiang Y, Liu N, Wang Q, et al. Endocrine Regulator rFGF21 (Recombinant Human Fibroblast Growth Factor 21) Improves Neurological Outcomes Following Focal Ischemic Stroke of Type 2 Diabetes Mellitus Male Mice. Stroke. 2018;49(12):3039–3049. doi:10.1161/STROKEAHA.118.022119

80. Sørensen G, Caine SB, Thomsen M. Effects of the GLP-1 Agonist Exendin-4 on Intravenous Ethanol Self-Administration in Mice. Alcohol Clin Exp Res. 2016;40(10):2247–2252. doi:10.1111/ACER.13199

81. Vallöf D, MacCioni P, Colombo G, et al. The glucagon-like peptide 1 receptor agonist liraglutide attenuates the reinforcing properties of alcohol in rodents. Addict Biol.2016;21(2):422–437. doi:10.1111/ADB.12295

82. Flippo KH, Trammell SAJ, Gillum MP, et al. FGF21 suppresses alcohol consumption through an amygdalo-striatal circuit. Cell Metab. 2022;34(2):317–328.e6. doi:10.1016/J.CMET.2021.12.024

83. Capozzi ME, Wait JB, Koech J, et al. Glucagon lowers glycemia when β-cells are active. JCI Insight. 2019;15(16). doi:10.1172/JCI.INSIGHT.129954

84. Hill CM, Laeger T, Dehner M, et al. FGF21 Signals Protein Status to the Brain and Adaptively Regulates Food Choice and Metabolism. Cell Rep. 2019;27(10):2934–2947.e3. doi:10.1016/J.CELREP.2019.05.022

